# Belowground allocation and dynamics of recently fixed plant carbon in a California annual grassland soil

**DOI:** 10.1101/2021.08.23.457405

**Authors:** Christina Fossum, Katerina Estera-Molina, Mengting Yuan, Don Herman, Ilexis Chu-Jacoby, Peter Nico, Keith Morrison, Jennifer Pett-Ridge, Mary Firestone

**Affiliations:** Department of Environmental Science, Policy, and Management, University of California, Berkeley, CA; Earth Sciences Division, Lawrence Berkeley National Laboratory, Berkeley CA; Physical and Life Sciences Directorate, Lawrence Livermore National Laboratory, Livermore, CA; Life and Environmental Sciences Department, University of California, Merced, CA

**Author notes:** Corresponding author: Mary Firestone, Department of Environmental Science, Policy, and Management, 140 Mulford Hall, University of California, Berkeley CA 94720.

**Keywords:** soil organic matter, annual grassland, ^13^CO_2_ pulse labeling, SOM density fractionation, ^13^C-NMR

## Abstract

Plant roots and the organisms that surround them are a primary source for stabilized organic C, particularly in grassland soils, which have a large capacity to store organic carbon belowground. To quantify the flow and fate of plant fixed carbon (C) in a Northern California annual grassland, we tracked plant carbon from a five-day ^13^CO_2_ pulse field labeling for the following two years. Soil and plant samples were collected immediately after the pulse labeling, and again at three days, four weeks, six months, one year, and two years. Soil organic matter was fractionated using a sodium polytungstate density gradient to separate the free-light fraction (FLF), occluded-light fraction (OLF), and heavy fraction (HF). Using isotope ratio mass spectrometry, we measured ^13^C enrichment and total C content for plant shoots, roots, soil, soil dissolved organic carbon (DOC), and the FLF, OLF, and HF. The HF was further analyzed by solid state ^13^C NMR spectroscopy.

At the end of the labeling period, the largest amount of ^13^C was recovered in plant shoots (60%), but a substantial amount (40%) was already found belowground in roots, soil, and soil DOC. Density fractionation of 4-week soil samples (from which living roots were removed) indicated that the highest isotope enrichment was in the mineral-rich heavy fraction, with similar enrichment of the FLF and OLF. At the 6-month sampling, after the dry summer period during which plants senesced and died, the amount of label in the FLF increased such that it was equal to that in the HF. By the 1-year sampling, ^13^C in the FLF had declined substantially and continued to decline by the 2-year sampling. ^13^C recovery in the OLF and HF, however, was qualitatively stable between sampling times. By the end of the 2-year experiment, 69% of remaining label was in the HF, 18% in the FLF and 13% in the OLF.

While the total ^13^C content of the HF did not change significantly from the 4-week to the 2-year sample time, ^13^C NMR spectroscopic analysis of spring HF samples from 2018, 2019, and 2020 suggests that the relative proportion of aliphatic/alkyl functional groups declined in the newly formed SOC over the 2-year period. Simultaneously, aromatic and carbonyl functional groups increased, and the proportion of carbohydrate groups remained relatively constant. In summary, our results indicate that initial associations between minerals and root-derived organic matter are significant and form rapidly; by 4 weeks, a substantial amount (17%) of the total plant-derived ^13^C had become associated with the heavy fraction (HF) of soil. While the majority of annual C input cycles rapidly (<2-year timescale), a sizeable proportion (∼12% of the original inputs) persisted for 2 years.

## 1. Introduction

California annual grasslands occupy over 10 million ha (Heady et al., 1992), and have been predicted to be a more resilient and therefore effective carbon sink than California’s fire-prone forests (Dass et al., 2018). According to a recent metaanalysis, grassland soils also have an increased capacity to store soil carbon in the face of increasing atmospheric CO_2_ concentrations (Terrer-Moreno et al., 2020). As such, these ecosystems have become an attractive target for California state policy initiatives aimed at mitigating atmospheric CO_2_ levels through soil carbon sequestration (Biggs & Huntsinger, 2021; Baker et al., 2020). However, our scientific understanding of soil carbon cycling, the fate of plant residues belowground, and how annual grasslands can broadly act as a viable carbon sink remains incomplete (Bradford et al., 2019). To effectively manage carbon stocks, further research is required to better understand the ecological parameters that regulate soil carbon cycling in these globally significant ecosystems.

Belowground carbon dynamics in California annual grasslands reflect the annual plant life cycle (Eviner & Firestone, 2007). Plant communities are dominated by annual grasses and forbs, species whose reproductive strategies are well-adapted to a Mediterranean climate where seasonal moisture limitations largely regulate nutrient cycling. This climate is characterized by cool wet winters and a warm summer period with no rainfall, a defining feature of which is the misalignment of two critical conditions for plant growth: rainfall and solar energy. Moisture limits plant growth in the summers, and sunlight and temperature limit plant growth in the winters (Bartolome et al., 2007; Eviner and Firestone, 2007). The plant life cycle begins with the first germinating rainfall in the autumn, proceeds slowly through the winter due to sunlight and temperature limitations, peaks in the spring when both soil moisture and sunlight are usually optimized, and ends with plant senescence once soil moisture has declined with the onset of the summer dry period (Eviner 2016; Eviner and Firestone, 2007). Throughout the winter/spring growing season, carbon enters the soil primarily via a constant “drip” of plant root exudation (Sokol & Bradford, 2019; Pett-Ridge et al., 2021); over the summer dry period, plant senescence and subsequent litter input, coupled with low-moisture conditions drives the transient accumulation of litter; in the autumn at the onset of the rainy season, a large portion of accumulated carbon is rapidly released as CO_2_ in a decomposition “pulse” (Blankinship & Schimel, 2018).

The characteristic seasonal soil moisture variability in Mediterranean climates can impact the persistence of soil organic matter (SOM) via multiple mechanisms. Following 4-6 months of no precipitation, autumn’s first major rainfall event strongly impacts the soil biota, physically stresses soil aggregates, chemically alters soil minerals, and destabilizes SOM, driving a pulse of respiration known as the Birch Effect (Barnard et al., 2020; Blazewicz et al., 2020; Birch, 1958). This release of CO_2_ is generally understood to result from the stimulation of microbial respiration and high availability of substrate (Schimel, 2018; Barnard et al., 2020). Plant germination usually occurs in the Autumn, temperature limits growth in the winter, and the peak period of plant growth occurs in the spring when both temperature and moisture are optimized. The majority of rhizosphere-derived SOC generation occurs during this spring growth period. Then, in late spring when soil moisture conditions no longer support growth, new root C inputs decline as plant senescence occurs and annual species set seed. During this dry-down period, decreasing soil moisture reduces hydrological connectivity between soil pores, isolating microbes from carbon substrates and resulting in a decline in ecosystem activity (Manzoni and Katul, 2014; Barnard et al., 2013; Blankinship and Schimel, 2018). Decomposition of plant biomass and transfer of carbon belowground continue to be limited throughout the summer dry period (Chou et al., 2008; Eviner and Firestone, 2007). While not fully understood, these wet-dry cycles characteristic of California annual grasslands may be important drivers of SOM persistence, and may be susceptible to future climate conditions that either amplify or minimize their stabilizing or destabilizing effects (Bailey et al., 2019).

Soil carbon is often considered in terms of distinct soil organic matter “pools”: either associated with minerals, occluded within microaggregates, or “free” (not subject to either chemical or physical protection) (Poeplau et al., 2018). Recent studies have shown that labile carbon substrates can play an important role in SOM formation and stabilization in both physically (occluded) and chemically (mineral-associated) pools (Cotrufo et al., 2015; Totsche et al., 2017; Villarino et al., 2021), while plant litter is generally understood to largely comprise the free-light fraction material. The mineral-associated or “heavy fraction” is of particular interest for soil organic carbon persistence; it is typically the oldest distinct pool (Torn et al., 1997) and represents carbon stabilized via mineral sorption and co-precipitation mechanisms (Kogel-Knabner et al., 2008). This fraction of SOM, commonly termed MAOM (mineral-associated organic matter) (Cotrufo et al., 2019), is largely of microbial origin and thought to be derived from relatively labile C substrates (Clemente et al., 2011; Cotrufo et al., 2013; Villarino et al., 2021). Recent work has suggested that microbial necromass may be a primary precursor to stable organic matter in grasslands (Angst et al., 2021).

In this study, we followed plant-fixed C entering the soil and moving into various soil organic matter pools, and also tracked its form and transformations over the course of multiple growing seasons. Initially, we exposed soil to the stable isotope ^13^C, via a 5-day ^13^CO_2_ field labeling of a California annual grassland plant community. To quantify the distribution and two-year dynamics of added ^13^C tracer in aboveground plant biomass, plant roots, soil, and various soil organic matter pools, we sampled biomass and soil at multiple timepoints following the field pulse labeling (immediately, 3 days, 4 weeks, 6 months, 1 year, and 2 years), and physically fractionated soil samples with a sodium polytungstate density gradient procedure. We isolated 3 soil density fractions: (HF) the heavy fraction (mineral-associated soil organic matter, MAOM), (OLF) the occluded-light fraction (micro-aggregate occluded soil organic matter), and (FLF) the free-light fraction (accessible soil organic matter debris) (Golchin et al., 1994; Sollins et al., 2009). We further characterized the chemistry of the heavy fraction ^13^C using solid state CPMAS ^13^C NMR spectroscopy, in order to better understand both the composition of carbon newly incorporated into this fraction, as well as the influence of new inputs on the total mineral-associated carbon pool. By combining ^13^C labeling, soil density fractionation, and solid-state ^13^C NMR, we sought to temporally characterize the flow and fate of newly fixed C entering this annual grassland soil and understand the dynamics of plant-derived soil organic carbon formation and persistence.

## 2. Methods

### 2.1 Field site description

Field work was conducted at the University of California Hopland Research and Extension Center (HREC), in southwestern Mendocino County, CA (39.004056, -123.085872). Climatic conditions at the field site are similar to conditions across Mediterranean California, with cool and wet winters, and hot, dry summers. Germination of the annual plant community occurs in the fall, following the first significant rainfall of the winter rainy season. Growth is limited throughout much of the winter by sunlight and temperature, and then peaks in the spring, followed by seed production and senescence by early summer (Becchetti et al., 2016). The vegetation community is dominated by naturalized annual grass and forb species including *Avena spp*., *Festuca spp.*, *Erodium spp.*, and *Bromus spp.* (Bartolome et al., 2007). Today, HREC is operated by the UC system as a working sheep ranch. Our field plots have been fenced off from grazing for >20 years. The soil at our field site belongs to the Squawrock-Witherell complex, a loamy-skeletal, mixed, superactive, thermic Typic Haploxeralf. Underlying parent material is colluvium derived from sandstone (Soil Survey Staff, 2020). Our measurements suggest the dominant clays are muscovite, chlorite, and kaolinite; dominant non-clay minerals are quartz and plagioclase (Table 1).

**Table 1.** Site characteristics and soil physicochemical properties. Location and soil pedological information determined via NRCS Web Soil Survey. Elevation and climate characteristics described by HREC. Soil texture and CEC determined by UC Davis analytical lab. Mineralogy determined at LLNL. Aggregate stability defined as % water-stable aggregates, was measured on soil collected March 2018. DOC, SOC, TN, C/N was collected in Spring 2018; pH and soil moisture was sampled in Spring 2018 during the rainy season and fall 2018 during the dry season.

|  |  |
| --- | --- |
| **Location** | 39°00'14.6"N 123°05'09.1"W |
| **Elevation (m)** | 244 |
| **MAT (°C)** | 14 |
| **MAP (cm)** | 94 |
| **Soil type** | Typic Haploxeralf |
| **Series name** | Squawrock-Witherell Complex |
| **Bulk density** | 1.457 g/cm<sup>3</sup> |
| **Aggregate stability** | 39 +/- 7 |
| **CEC (meq/100g)** | 18.75 |
| **SOC (mg/g soil)** | 13.1 +/- 1.7 |
| **TN (mg/g soil)** | 1.3 +/- .1 |
| **C/N** | 9.7 +/- .4 |
| **Texture** | 48, 35, 17 |
| **Clays (30%)** | Muscovite (17.8%) Chlorite (11.8%)<br>Kaolinite (0.7%) |
| **Non-clays (70%)** | Quartz (47.3%) Plagioclase<br>(22.4%) |
| **pH (Spring)** | 5.9 +/- .1 |
| **pH (Fall)** | 7.3 +/- .1 |
| **Soil Moisture % (Spring)** | 14.3 +/- 2.6 |
| **Soil Moisture % (Fall)** | 1.9 +/- .3 |

### 2.2 Experimental design

Samples collected for this study are a subset from a larger ^13^CO_2_ labeling and precipitation manipulation field experiment. In the spring of 2017, sixteen 3.24 m^2^ plots were established, each delineated by a 1-m deep plastic liner to limit soil water equilibration with surrounding soils, and each containing six 40-cm diameter circular subplots. Circular subplots were surrounded by a 15cm deep PVC “collar” designed to be fitted with an above-ground cylindrical chamber for labelling plants with either ^12^C or ^13^C-CO_2_ (Supplemental Figure 1A). Each circular subplot was subdivided into four 15-cm deep sections via plexiglass dividers; this “wedge” design allowed us to destructively harvest ^13^CO2-labeled soil and biomass from a single circular subplot at multiple timepoints following a labeling event (Supplemental Figure 1B). Removable rain-exclusion shelters were installed above all plots and precipitation manipulation began in the fall of 2017 and continued through the 2019-2020 growing season. Of the 16 plots, eight received 50% of average annual precipitation and eight received 100%, where average annual rainfall was calculated from precipitation data collected at HREC dating back to 1951 (Supplemental Figure 2). Depending on the timing and quantity of natural rainfall events, field plot precipitation was augmented via manual watering with natural spring water and limited through application of the rain-out shelters.

Experimental plots were labeled with ^12^C or ^13^C-CO_2_ (99 atm %, Cambridge Isotopes) for 5 days from February 11-15, 2018, timed to correspond with the expected maximum root development phase of *Avena spp.* plant growth (the plots were seeded the prior year to encourage a *Avena spp.* dominated community). CO_2_ levels during the pulse labeling event were monitored with a Picaroon G2200-I analyzer (for ^13^C) and infrared gas analyzer (IRGA) (for ^12^C). For the labeling, well-sealed cylindrical chambers made of PAR transmissive PVC film were fitted over two circular subplots per plot (Supplemental Figure 1A). Plants within these chambers were exposed to either ^12^CO_2_ or an isotopically labeled analog, ^13^CO_2_, resulting in 16 ^12^CO_2_-labeled ‘control’ subplots and 16 ^13^CO_2_-labeled subplots. The headspace CO_2_ concentrations within the chambers were maintained between 400-1500ppm during the daytime, and chambers were removed from plots at night. ^13^C enrichment of ^13^CO_2_-labeled subplots was maintained between 30-75 atom%. Within each CO_2_ group, 8 of the 16 subplots were under the 50% precipitation regimen, and 8 under the 100% regimen.

### 2.3 Field harvests of ^13^C labeled biomass and soil

Soil was harvested at six times following the ^13^CO_2_ labeling event: (1) immediately after the 5-day labeling period, (2) three days later, (3) four weeks after the first harvest, (4) six months after the first harvest, (5) one year after the first harvest, and (6) two years after the first harvest. We subsequently refer to these harvest sampling times both by the time elapsed since the ^13^C labeling event (0 days, 3 days, 4 weeks, 6 months, 1 year, and 2 years), as well as by the season in which the harvest occurred: Spring ‘18 (4 weeks), Fall ‘18, Spring ‘19, and Spring ‘20. At each sampling time, we harvested 4 replicate samples from each experimental treatment: ^12^C-labeled, ^13^C-labeled, 50% precipitation, 100% precipitation. We did not detect a statistically significant (p < 0.05) precipitation treatment effect on any soil or plant characteristics measured in this study; thus, replicates from these treatments were pooled for final analyses such that n=8 for most of the analyses presented.

Root and shoot biomass samples were collected at three times following the ^13^CO_2_ pulse labeling: (1) at zero days, when we expected the maximum ^13^C-labeled shoot biomass, (2) at three days, and (3) at four weeks, by which point we expected a measurable fraction of ^13^C label to have been translocated to root biomass and the surrounding soil. Shoot biomass was collected by clipping all live aboveground plant tissue from a single subplot “wedge”, and then scaling up to g / m^2^. Root biomass was collected from three 2.45-cm cores that were installed in each wedge at the time of harvest and scaling up to the full wedge volume (1/4 X 40cm subplot X 15cm depth), and then g / m^2^. Prior to the 6-month sampling time (Fall18), aboveground litter remaining from the 2017-2018 growing season was removed from the plots.

At each of the first four sampling times, we harvested 16 “wedges” of soil. Once soil wedges were harvested, visible roots were removed and soil was homogenized, 500g sub-samples were collected, air-dried, and stored at ambient temperature for further analysis. At each of the final two sampling times (Spring19 and Spring20), we harvested two cores per wedge (2.54 cm diameter X 15cm deep) rather than the whole wedge to preserve the remaining soil for future experiments, however, we harvested the same number of samples as in the previous two harvests. In these two cores we removed roots and homogenized the soil (∼300g) as above, then air dried and stored the soil for further analyses. Additional subsamples at each of the four harvests were immediately aliquoted from homogenized soil for analyses requiring fresh soil.

Altogether, eight ^13^C-labeled and eight ^12^C-labeled shoot, root, and soil samples were harvested at 0 days, 3 days, and 4 weeks after labeling. Senescence of ^13^C-labeled plants occurred over the summer of 2018, between the 4-week and 6-month sampling times. Therefore, for the remaining sampling times (6-month, 1-year, and 2-year), only soil samples were collected (eight ^13^C-labeled and eight ^12^C-labeled).

### 2.4 Processing and diagnostic analyses of soil, root, and shoot biomass

Aboveground biomass dry weight was determined by drying harvested biomass at 65 °C until a stable dried weight was achieved. Root biomass dry weight was determined by hand picking live roots from collected root biomass cores, washing, and drying at 65 °C until dried weights reached a stable plateau. Dry root biomass per wedge was then calculated by scaling the dried root weights to the volume of the wedge.

Soil pH was determined in a 1:1 ratio of fresh homogenized soil to 0.01M CaCl_2_. Soil gravimetric water content was assessed by drying a 10g subsample of fresh homogenized soil at 105 °C until dried weights reached a stable plateau when weighed, then calculating percent water. DOC was extracted from 5g fresh soil with 20ml 0.5M K_2_SO_4_; DOC and ^13^C-DOC was assessed by the Yale Analytical and Stable Isotope Center (YASIC) via wet oxidation method (Lang et al., 2012). Remaining analyses were conducted on homogenized air-dry soil. Soil texture analysis was conducted by the UC Davis Analytical Lab via the hydrometer method (Sheldrick et al., 1993). Aggregate stability was conducted using a wet sieving method previously adapted by Sher et al. (2020), using a custom wet-sieve apparatus (Singer et al., 1992; Sher et al., 2020).

### 2.5 Soil quantitative X-ray diffraction analysis

Soil mineralogy was determined at Lawrence Livermore National Lab (Zhou et al., 2018). Soil samples were dried, crushed and passed through a 500 µm sieve. Then 3 grams of soil was ground with 15 mL of methanol in a McCrone mill with corundum grinding elements for 5 minutes. The sample was transferred into a plastic tray, air dried and homogenized on a vortex mixer with 10 mm plastic beads for 3 minutes (Bakker et al., 2018). The random powders were then side loaded into XRD sample holders and analyzed on a Bruker D8 advance XRD scanning from 3 to 65° 2Θ with a step size of 0.011° at a rate of 5 seconds per step. Quantitative analysis was done using BGMN Rietveld refinement and the Profex interface software (Doebelin et al., 2015). The XRD patterns were refined to fit crystal unit cell parameters, size, site occupancy and preferred orientation.

### 2.6 Soil density fractionation

Soil density fractionation was performed on samples collected at 4-weeks, 6-months, 1-year, and 2-year sampling times. Air-dried soil was sieved to 2mm before being density fractionated into three discrete pools of soil organic matter using a sodium polytungstate (SPT) density gradient: free-light fraction (FLF, ρ < 1.75g-cm^-3^), occluded-light fraction (OLF, ρ < 1.75g-cm^-3^), and mineral-associated or heavy fraction (HF, ρ > 1.75g-cm^3^). The density of 1.75g-cm^-3^ was chosen due to the similarities in mineralogy and soil physical characteristics between this sampling site and the site sampled in *Neurath et al., 2021*, which used this same SPT approach.

The method for density fractionation used in our study was adapted from Hicks Pries et al. (2017), previously adapted from Strickland & Sollins (1987). For each sample, 50mL of sodium polytungstate (SPT-0, Geoliquids) prepared to a density of 1.75g-cm^-3^ was added to a 250mL centrifuge tube containing 20g of air-dried soil. The mixture was inverted by hand to ensure all soil came into contact with the SPT; soil remaining on the lid and sides was rinsed down with an additional 50mL of SPT. The SPT-soil solutions were allowed to settle for 1 hour, then centrifuged for 1 hour at 3,700 RCF in a swinging bucket rotor (Beckman Avant J-20 Floor Centrifuge with JS-5.3 rotor). Following centrifugation, samples were allowed to settle again until no particles remained suspended within the SPT solution. Particles floating on top of the SPT solution were defined as the FLF and were isolated by aspirating onto a 0.7μm glass microfiber filter (Wattman), and rinsed with MilliQ water to remove residual SPT. Then, the FLF filters were transferred into drying tins in a 55 °C oven until standing water had evaporated.

To release the OLF from microaggregates, the remainder of the soil-SPT mixture was mixed with a benchtop mixer for 1 minute, followed by sonication for 90 seconds. As above, the soil was then rinsed, allowed to settle, centrifuged for 1 hour, and the floating fraction isolated via aspiration and dried.

The remaining sediment (ρ > 1.75g-cm^-3^) was defined as the HF. This fraction was rinsed with 150mL Milli-Q H2O, vigorously shaken by hand, centrifuged for 20 minutes, followed by aspiration and disposal of the supernatant. This was repeated 5 times, or until the density of the supernatant ∼ 1g-cm^-3^. The HF was transferred into drying tins and dried at 55 °C. Once standing water had evaporated, all three fractions were transferred into a 105 °C oven for 48 hours. Oven-dried samples were cooled in a desiccator before being weighed, ground with a mortar and pestle, and stored in glass vials.

### 2.7 Isotopic and elemental analysis

Prior to elemental analysis, samples were ground to a fine powder and weighed into aluminum tins, with sample weight proportional to expected carbon content (4mf for the FLF, 2mg for the OLF, 50mg for the HF and bulk soil, 0.35mg for shoot biomass, and 0.30mg for root biomass samples). Bulk and density fractionated soil were ground using a mortar and pestle, aboveground biomass was ground using a coffee grinder, and root biomass was ground by hand. Total C, N, and ^13^C enrichment were measured via total combustion using an elemental analyzer coupled with a continuous flow Isotope Ratio Mass Spectrometer (IRMS) at the Stable Isotope Facility at the University of California, Davis (AOAC Official Method 972.43).

### 2.8 ^13^C-NMR

Subsamples of heavy fraction (HF) material were analyzed by ^13^C-NMR to assess broad chemical composition of mineral-associated organic matter. Solid-state ^13^C cross polarization magic angle spinning (CPMAS) spectroscopy was run on one ^13^C-labeled HF sub-sample from the 4-week, 1-year, and 2-year sampling times, as well as one ^12^C-labeled control HF subsample from the 4-week timepoint as a control comparison. These four spectra were acquired using a 4mm log-gamma CPMAS probe on a 500 MHz Bruker Avance 1 NMR spectrometer at the UC Davis NMR facility. Samples were spun at 10 kHz with an acquisition time of 41 ms. Scan number ranged from 75,000 – 102,400. Glycine (176ppm) was used as the external reference. Using Topspin software, data were zero filled to 8k; an exponential function with 500 Hz of line broadening was used for signal processing, with zero order phase correction, followed by manual baseline correction.

Broad C functional groups were defined based on the following chemical shift regions: aliphatic/alkyl C (0-45ppm), O-alkyl C (45-110ppm), aromatic and aryl C (110-162ppm), and carbonyl C (162-190ppm) (Helfrich, 2006). Integration of chemical shift regions was conducted using Topspin 3 software to calculate relative contribution of different functional group regions to total peak area (0-190ppm).

### 2.9 Statistical analyses

Effect of sampling time on C, N, and ^13^C content was determined using ANOVA. Statistical significance was determined using Tukey’s HSD post-hoc test with the R package ‘agricolae’ (Mendiburu, 2015). Data was visualized using the R package ‘ggplot2’ (Wickham, 2016).

### 2.10 Calculation of ecosystem ^13^C assimilation

We define ecosystem assimilated ^13^C as that quantifiable in (1) aboveground (plant biomass) and (2) belowground (soil + root biomass) pools, in other words net ^13^C gain rather than gross ecosystem exchange. Ecosystem ^13^C assimilation was calculated by converting the ^13^C concentration of each pool (aboveground biomass, root biomass, soil in mg / g) to quantity (g ^13^C / m^2^ over a 15cm sampling depth) and then summing. For simplicity, the sum of ^13^C recovered in aboveground biomass plus soil and root biomass immediately after the pulse labeling period (0-days) was interpreted to equal 100% of assimilated ^13^C. The percentage of assimilated ^13^C remaining at each subsequent sampling timepoint was then calculated relative to this original amount (Table 2). We note that, at the 6-month, 1-year and 2-year sampling times, ^13^C-labeled aboveground biomass and roots had senesced and the aboveground pool was not measured, and roots were not physically separately from the soil as was done for the 0-days and 4-weeks timepoints. Average ^13^C recovery based on summing the FLF + OLF + HF was approximately 72% that of the bulk soil, which is within the range for density fractionated soil C recovery values cited in the literature (Crow et al., 2007; Cusack et al., 2018).

**Table 2:**
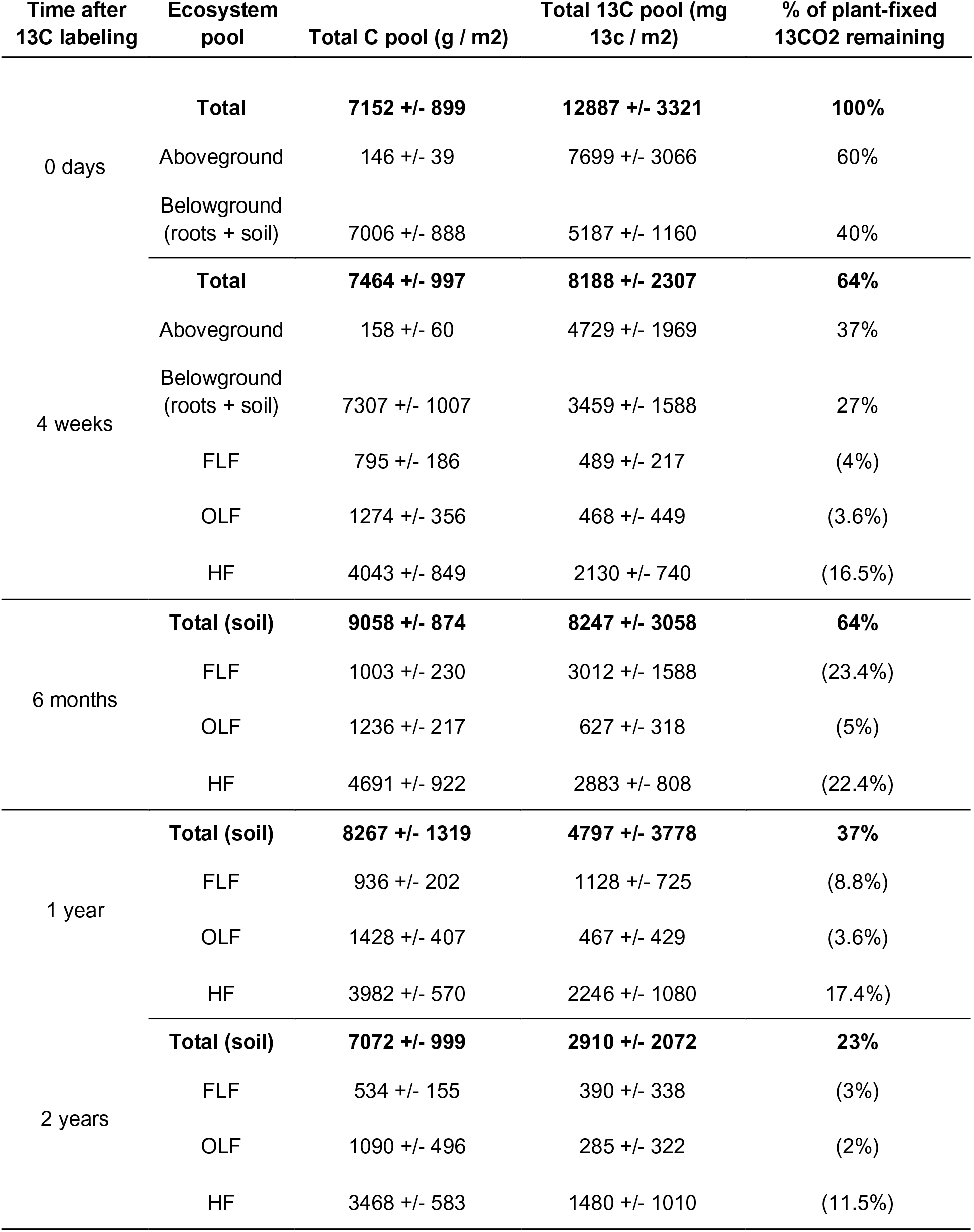
Distribution of ^13^C label within ecosystem pools. We defined 100% remaining plant-fixed ^13^CO_2_ as that present in the system (aboveground + belowground pools) immediately after the ^13^CO_2_ labeling period (“0 days” timepoint). The aboveground pool consists of aboveground plant biomass, and the belowground pool consists of root biomass + bulk soil. For the “6 months”, “1 year” and “2 year” timepoints, ^13^C-labeled roots had senesced and so were not measured separately from the bulk soil was done for the “0 days” and “4 weeks” timepoints. The bulk soil was further separated into three sub-pools via soil density fractionation, yielding the free-light fraction (FLF), occluded-light fraction (OLF), and heavy fraction (HF). Average ^13^C recovery based on FLF + OLF + HF was about 72% that of the bulk soil, which is within the range for C recovery in density fractionated soil generally cited in the literature (Crow et al., 2007; Cusack et al., 2018).

## 3. Results

### 3.1 Ecosystem ^13^C Incorporation

Immediately following the ^13^CO_2_ labeling period, all four ecosystem pools that we analyzed (soil, shoot and root biomass, DOC) were enriched in ^13^C (Figures 1, 2). We defined the total amount of ^13^C present at the 0-day to be 100% and calculated pools thereafter relative to this starting point. During the 5-day labeling period, over 12g ^13^C / m^2^ derived from plant photosynthate had accumulated in these ecosystem pools, accounting for roughly 0.2% of total ecosystem C content (aboveground biomass, root biomass, and soil in the 0-15cm depth horizon). By 4 weeks, shoot biomass ^13^C content had declined to 61.7% of the initial amount (p = 0.015) (Figure 1, Table 2). Transfer of plant-fixed ^13^C to belowground pools was immediate, accounting for 40% of total ecosystem ^13^C immediately after the labeling. Belowground ^13^C content reached a peak 6 months post-labeling. Between the 6-month, 1-year, and 2-year sampling times, assimilated ^13^C in the ecosystem decreased stepwise: 64% of the original ^13^C remained at 6 months, 37% at 1 year, and 23% at 2 years (Figure 1, Table 2).

**Figure 1.**
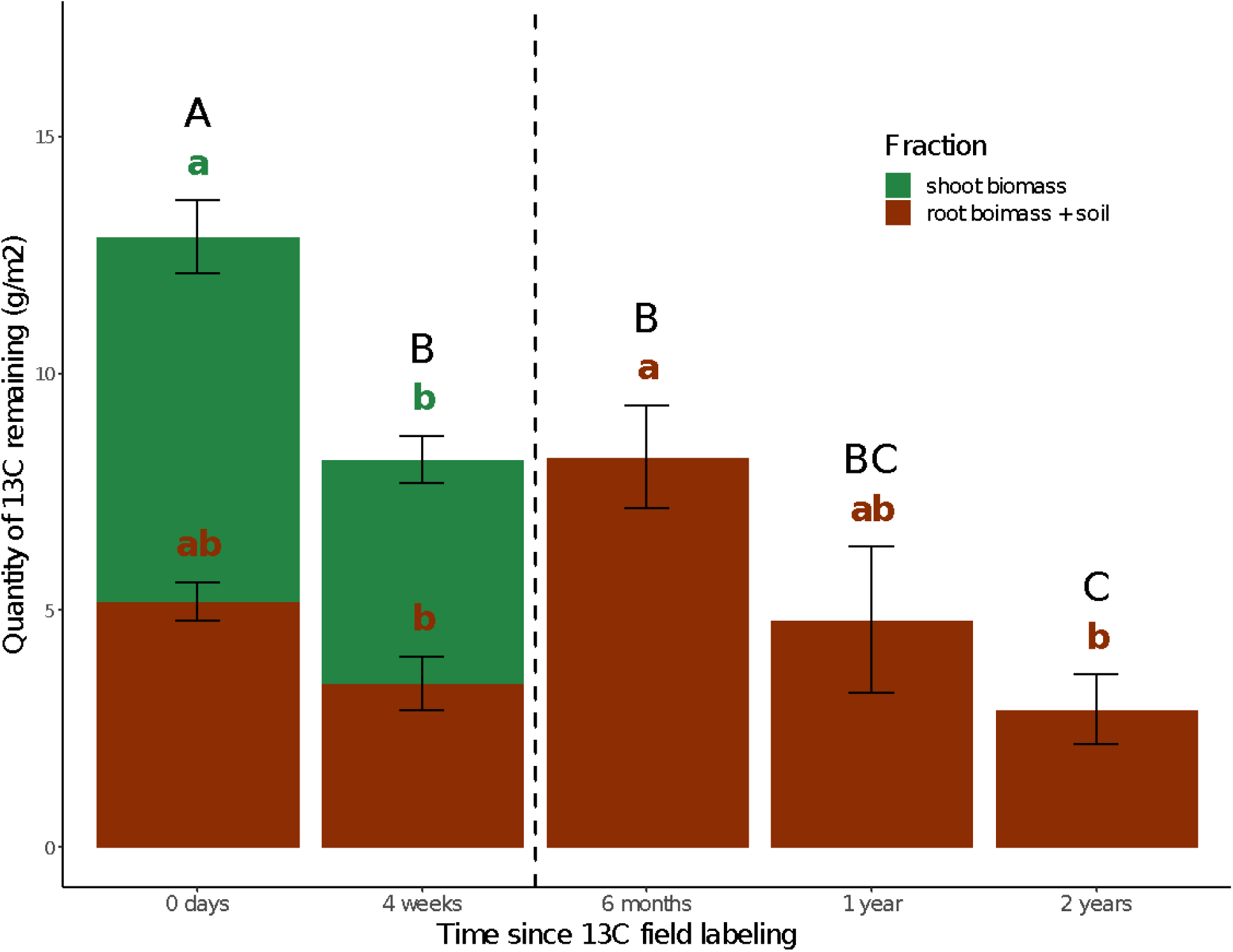
Ecosystem ^13^C assimilation. Total excess ^13^C measured in shoot biomass (green) and root biomass + soil (brown). The 0-day timepoint occurred immediately following completion of the 5-day ^13^CO_2_ labeling. Vertical black dashed line between the 4-weeks and 6-months timepoint marks plant senescence at the end of the spring growing season—hence, no shoot biomass ^13^C measurements occurred beyond that point and previously living root ^13^C biomass has become part of the soil ^13^C. Letters indicate significant differences within a given ecosystem fraction (shoots in green vs. root biomass + soil in brown text), and overall (in black text) between timepoints by Tukey’s HSD (p-value = 0.05). Error bars represent 1 SE.

**Figure 2.**
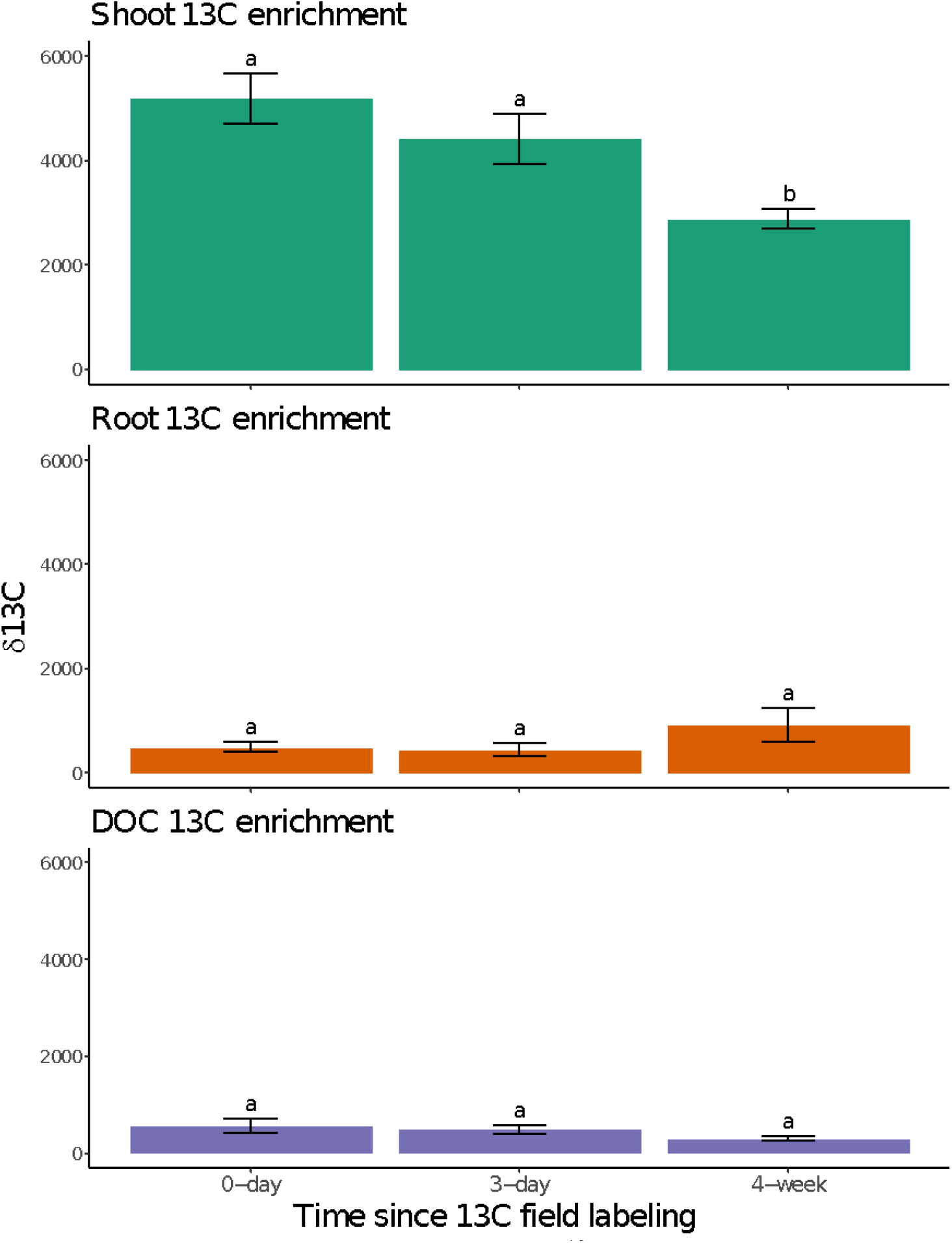
Short-term ecosystem ^13^C dynamics. Delta ^13^C values measured in shoot biomass (top), root biomass (middle), and soil DOC (bottom). Timepoints correspond to 0 days, 3 days, and 4 weeks following the completion of the 5-day ^13^CO_2_ field labeling. Letters indicate significant differences within a given ecosystem fraction (aboveground biomass, roots, DOC) between timepoints by Tukey’s HSD (p-value = 0.05). Error bars represent 1 SE.

Samples collected at 0 days, 3 days, and 4 weeks following ^13^CO_2_ labeling were used to assess the initial dynamics of ^13^C in shoots, roots and DOC (Figure 2). Shoot ^13^C enrichment (g ^13^C / m^2^) declined significantly over the 4 weeks post labeling. While not statistically significant at p < 0.05, mean shoot biomass ^13^C appeared to decline slightly by 3 days post-labeling, despite no detectable change in shoot biomass--possibly due to loss of recently fixed ^13^C via plant respiration. Allocation of ^13^C-labeled photosynthate to the roots occurred immediately, and did not significantly change over the 4 weeks, despite expected dilution of the atom-% ^13^C by continued root growth. ^13^C enrichment of the DOC pool did not significantly change between 0 days, 3 days, and 4 weeks following the ^13^CO_2_ labeling event.

### 3.2 Soil Density Fractions

We used density fractionation to assess changes in the quantity of newly-fixed plant-derived C in soil samples collected in March of 2018 (Spring18 = 4-week sampling time), the Fall of 2018 (Fall18 = 6-month), April of 2019 (Spring19 ∼ 1-year), and March of 2020 (Spring20 ∼ 2-year). Of the three fractions assessed, the HF accounted for roughly 99% of the total recovered soil mass (Table 3) and contained the highest amount of C, accounting for 66% of the total on average, while the OLF was 20%, and FLF 14% (Figure 3). The distribution of total N followed a similar pattern to total C, but with an even greater proportion of N accumulating in the HF (Table 3): 83% in the HF, 10% in the OLF, and 7% in the FLF. The C:N ratio varied significantly by fraction type: highest in FLF (∼20:1), intermediate in OLF (∼18:1), and lowest in HF (∼8:1), and these differences were statistically significant (p < 0.05). Total C, N, and C:N (Table 3, Figure 3) did not exhibit detectable seasonal fluctuations between the Spring and Fall sampling times. For all density fractions, total C was generally lower at the Spring20 sampling than at other sampling times (Figure 3).

**Figure 3.**
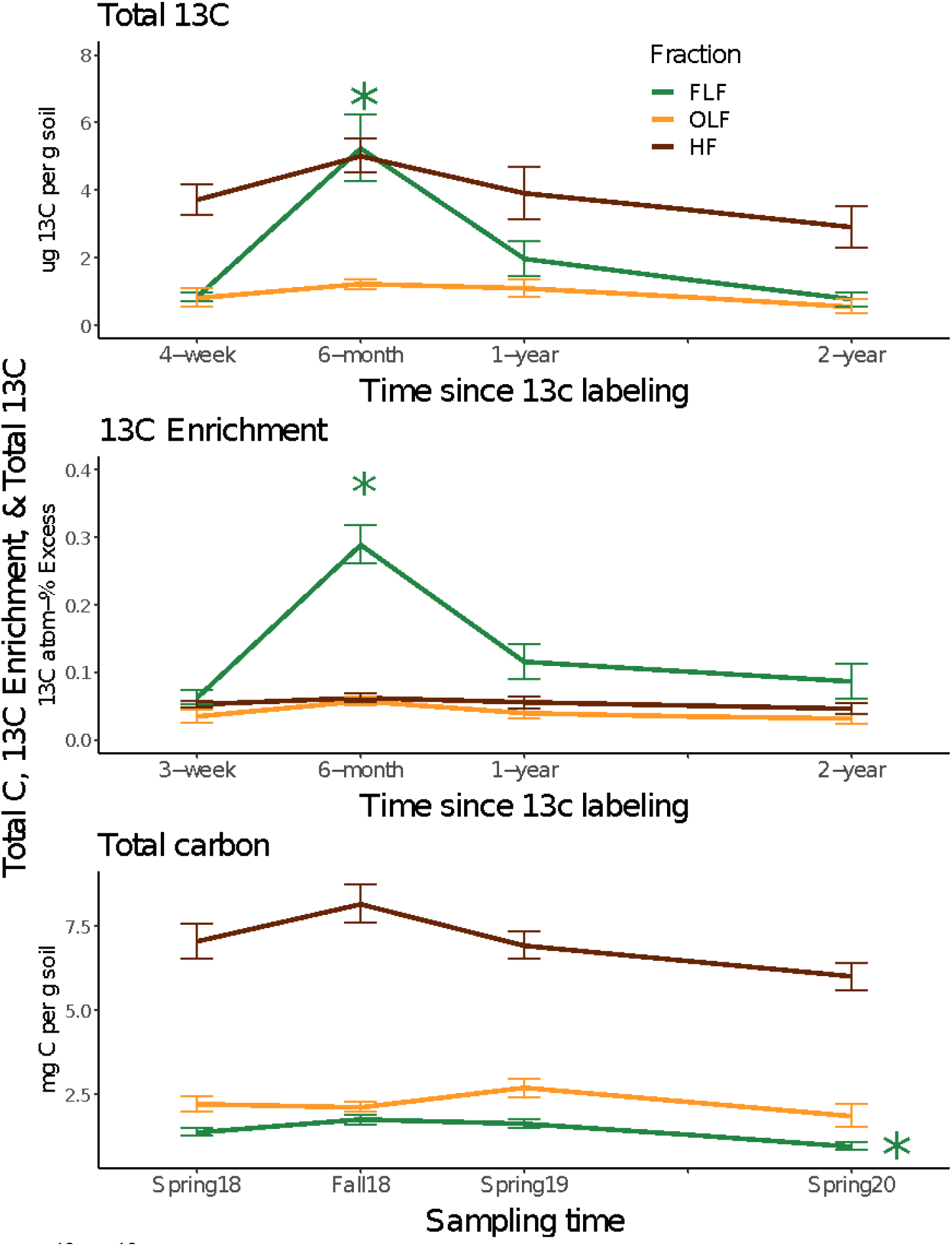
Total ^13^C, ^13^C enrichment, and total C distribution among soil density fractions. Total ^13^C (ug / g soil), ^13^C enrichment (atom-% excess), and total carbon (mg / g soil) measured for soil density fractions: free-light fraction (FLF), occluded-light fraction (OLF), and heavy fraction (HF). Soil density fractions were determined in Spring18, Fall18, Spring19, and Spring20. These sampling times correspond to 4-weeks, 6-months, 1-year, and 2-years following the ^13^CO_2_ field labeling. Error bars represent 1 SE. Significant differences by Tukey’s HSD (p-value = 0.05) between times are indicated by stars with color coordinating to respective soil density fraction.

**Table 3:**
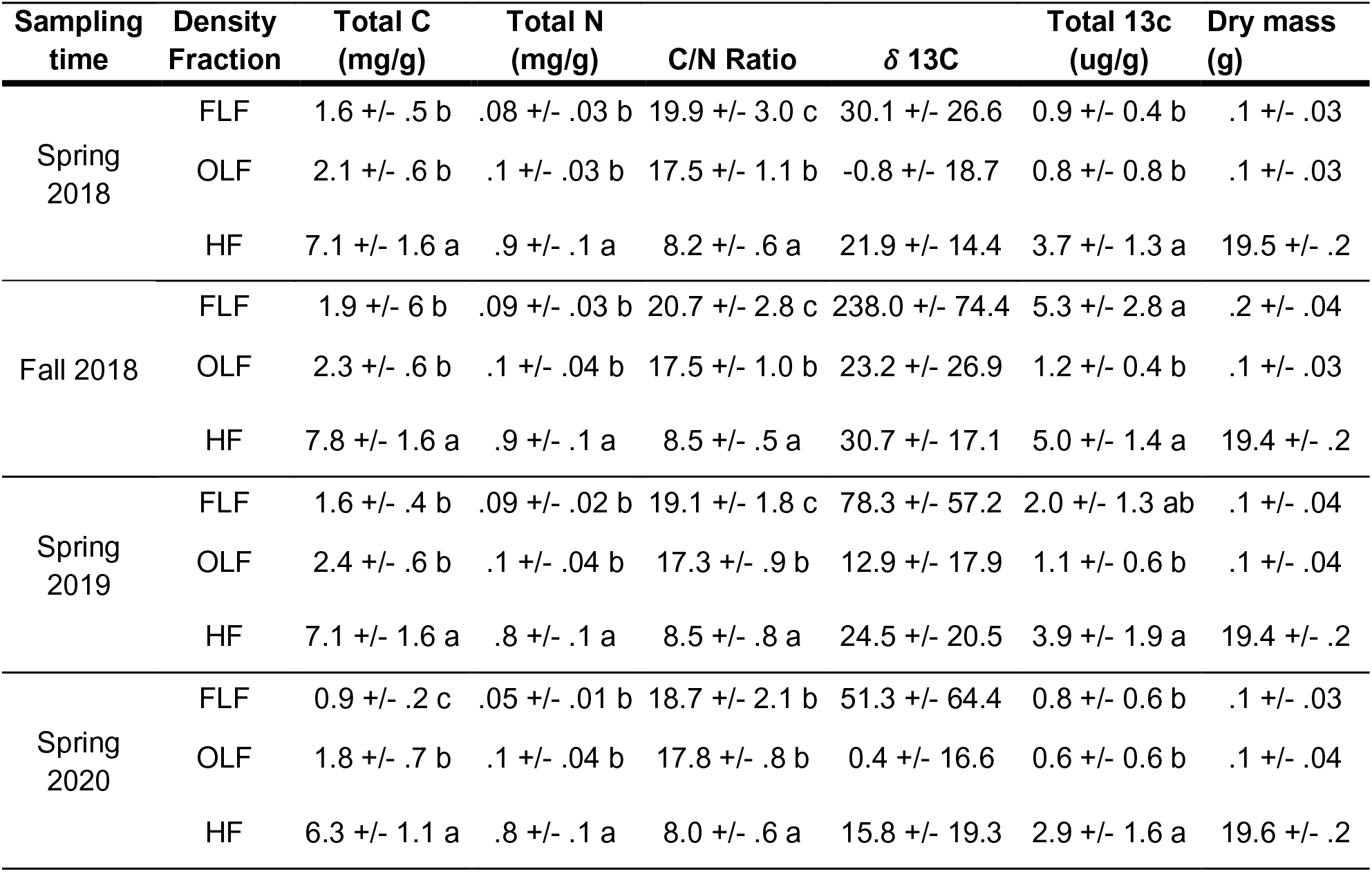
Characteristics of soil density fractions. Means shown +/- 1SD. Elemental characteristics are shown in units of quantity of element per gram of soil. δ^13^C values represent absolute ^13^C enrichment. Total ^13^C values represent total ^13^C added during the ^13^C labeling period (calculated from atom-% excess ^13^C). Dry mass (g) value represents quantity of starting soil sample (∼20g) recovered in each density fraction.

| Sampling time | Density Fraction | Total C (mg/g) | Total N (mg/g) | C/N Ratio | $\delta^{13}\text{C}$ | Total $^{13}\text{C}$ (ug/g) | Dry mass (g) |
| --- | --- | --- | --- | --- | --- | --- | --- |
| Spring 2018 | FLF | 1.6 +/- .5 b | .08 +/- .03 b | 19.9 +/- 3.0 c | 30.1 +/- 26.6 | 0.9 +/- 0.4 b | .1 +/- .03 |
|  | OLF | 2.1 +/- .6 b | .1 +/- .03 b | 17.5 +/- 1.1 b | -0.8 +/- 18.7 | 0.8 +/- 0.8 b | .1 +/- .03 |
|  | HF | 7.1 +/- 1.6 a | .9 +/- .1 a | 8.2 +/- .6 a | 21.9 +/- 14.4 | 3.7 +/- 1.3 a | 19.5 +/- .2 |
| Fall 2018 | FLF | 1.9 +/- .6 b | .09 +/- .03 b | 20.7 +/- 2.8 c | 238.0 +/- 74.4 | 5.3 +/- 2.8 a | .2 +/- .04 |
|  | OLF | 2.3 +/- .6 b | .1 +/- .04 b | 17.5 +/- 1.0 b | 23.2 +/- 26.9 | 1.2 +/- 0.4 b | .1 +/- .03 |
|  | HF | 7.8 +/- 1.6 a | .9 +/- .1 a | 8.5 +/- .5 a | 30.7 +/- 17.1 | 5.0 +/- 1.4 a | 19.4 +/- .2 |
| Spring 2019 | FLF | 1.6 +/- .4 b | .09 +/- .02 b | 19.1 +/- 1.8 c | 78.3 +/- 57.2 | 2.0 +/- 1.3 ab | .1 +/- .04 |
|  | OLF | 2.4 +/- .6 b | .1 +/- .04 b | 17.3 +/- .9 b | 12.9 +/- 17.9 | 1.1 +/- 0.6 b | .1 +/- .04 |
|  | HF | 7.1 +/- 1.6 a | .8 +/- .1 a | 8.5 +/- .8 a | 24.5 +/- 20.5 | 3.9 +/- 1.9 a | 19.4 +/- .2 |
| Spring 2020 | FLF | 0.9 +/- .2 c | .05 +/- .01 b | 18.7 +/- 2.1 b | 51.3 +/- 64.4 | 0.8 +/- 0.6 b | .1 +/- .03 |
|  | OLF | 1.8 +/- .7 b | .1 +/- .04 b | 17.8 +/- .8 b | 0.4 +/- 16.6 | 0.6 +/- 0.6 b | .1 +/- .04 |
|  | HF | 6.3 +/- 1.1 a | .8 +/- .1 a | 8.0 +/- .6 a | 15.8 +/- 19.3 | 2.9 +/- 1.6 a | 19.6 +/- .2 |

The ^13^C labeling event occurred during the growing season in February 2018, and samplings occurred in Spring 2018, Fall 2018, Spring 2019, and Spring 2020. Four weeks after the ^13^C labeling, 24% of the initial ecosystem ^13^C was recovered in soil density fractions (Table 2). Of that, significantly more was recovered in the HF than in the OLF or FLF. While accumulation of total C was higher in the OLF than FLF, ^13^C was in the FLF at the 4-week sample time. ^13^C enrichment (atom-% ^13^C) was generally highest in the FLF and lowest in the OLF. Between the 4-week and 6-month sampling times, soil organic ^13^C-labeled carbon (SO^13^C) recovered in the density fractions roughly doubled, with the majority of additional SO^13^C accumulating in the FLF (Figure 3, Table 2). At 6 months, the amount of ^13^C recovered in the FLF was similar to that in the HF. Between 6 months and 1 year after labeling, SO^13^C content of the density fractions declined by 40%, with a particularly large decrease observed in FLF material. The 2-year ^13^C recovery in the soil density fractions was 55% of that recovered in the 1-year samples. By Spring20, over two years after the original ^13^C addition, isotopically labeled carbon persisted in all three fractions, with over 50% of the remaining ecosystem ^13^C (representing 13.5% of the initial ecosystem ^13^C content) found in protected forms, either occluded within soil microaggregates (OLF) or associated with soil minerals (HF) (Table 2).

### 3.3 ^13^C NMR analysis of mineral-associated carbon

Heavy fraction (HF) material from 4-week, 1-year, and 2-year sampling times was analyzed by ^13^C NMR to assess the molecular forms taken by the newly fixed ^13^C and present in the HF. Likely due to the presence of paramagnetic minerals in these soils, the peaks in the ^13^C NMR spectra were broad and hence we were not able to identify specific compounds. Instead, we assessed relative proportions of broad chemical classes based on chemical shift regions in the NMR spectra: alkyl C 0-45ppm, O-alkyl C 45-110ppm, aromatic C 110-162ppm, and carbonyl C 162-190ppm (Figure 4, Supplemental Figure 5).

**Figure 4:**
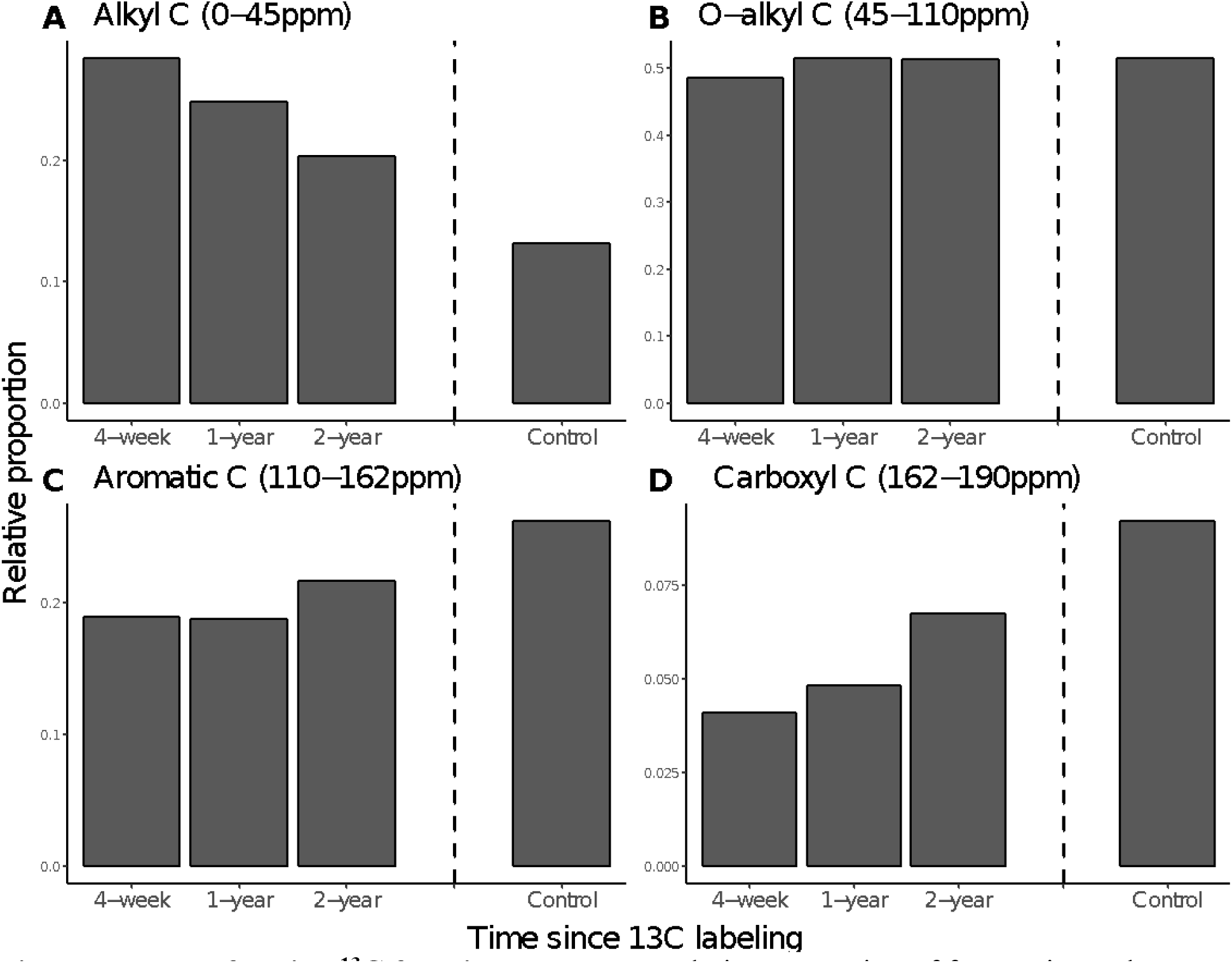
Heavy fraction ^13^C functional groups. Relative proportion of four major carbon functional groups in heavy fraction separated from soil calculated from ^13^C NMR spectra. Proportions were calculated by integrating functional group regions A. alkyl C, B. O-alkyl C, C. aromatic C, and D. carboxyl C using Topspin software, and analyzed separately for each spectrum. X axis refers to spectra obtained for ^13^C-labeled Heavy Fraction soil 4 weeks, 1 year, and 2 years following the ^13^C-labeling, as well as a ^12^C-labeled control sample collected in the Spring of 2018. Atom percent ^13^C for the 4 samples (4-wk, 1 year, 2 years, and control) was 1.145, 1.139, 1.133, and 1.076 respectively.

Labeling plants with ^13^CO_2_ allowed us to follow the functional group characteristics of newly fixed C incorporated into the mineral-associated pool (Figure 4). The SO^13^C-HF appeared to be relatively enriched in alkyl C with lesser amounts of aromatic and carboxyl C. In the ^13^C NMR spectra, the relative proportion of alkyl C declined over time from 4 weeks to 1 year to 2 years after the ^13^C labeling period. Over this same period, the relative proportion of carboxyl C increased, while the relative proportion of aromatic C and O-alkyl C remained constant. While the proportion of C functional groups in the ^13^C-labeled HF material was distinct from the ^12^C-labeled control HF material at all sampling times, the relative proportions of C functional groups in the ^13^C-labeled material appeared to become more similar to those observed in the control material over time.

## 4. Discussion

### 4.1 Ecosystem ^13^C Incorporation

In this California annual grassland soil, growing plants quickly allocated a substantial proportion of photosynthate belowground. We traced the translocation of plant photosynthate to belowground carbon pools during a 5-day ^13^CO_2_ field labeling (Figure 1). Our labeling period occurred in late winter (February 11-15, 2018), which was intended to correspond to the plant growth stage of maximum allocation of aboveground photosynthate C to root growth (Jackson et al., 1989). By the end of this 5-day period, 40% of assimilated ^13^C had been allocated belowground (Figure 1).

^13^C recovery in plant roots is a clear indication of the incorporation of fresh plant-derived carbon inputs into the soil, particularly as these ^13^C-labeled roots decompose over time. We observed evidence of this phenomenon at the 6-month sampling time, when ^13^C recovery in FLF material reached its peak. We presume this was primarily due to incorporation of ^13^C labeled root litter from the previous growing season. However, at the 4-week sampling time, we already observed high ^13^C recovery (nearly 38%) in all our soil density fractions, from which we had removed live roots. This suggests that substantial ^13^C was exuded into the soil by the roots during this period of plant growth, or that other types of rhizodeposits (sloughed root tip cells, cell hairs) had become part of the soil’s organic matter pools (Table 2).

In our study, the ^13^C enrichment of root tissue was similar to the enrichment of the DOC pool when measured either immediately or 3 days after the end of the labeling period (Figure 2). This suggests that root exudates were in equilibrium with root biomass enrichment during this plant growth stage. By the 4-week sampling time, ^13^C enrichment appears to slightly increase in plant roots and decline in soil DOC (although neither increase nor decrease was statistically significant), which could imply a shift of plant C allocation from labile root exudates, to root structural compounds. Furthermore, we observed a slight decline in belowground ^13^C content between 0 days and 4 weeks following the ^13^C labeling (Figure 1), which is likely from root and microbial respiration. These findings indicate that as plants continue to grow, recently-fixed photosynthate was rapidly translocated within the plant and released into the soil as labile C compounds exuded by roots (and perhaps also by associated arbuscular mycorrhizal fungi (AMF) (Kakouridis et al., 2021)). Almost just as rapidly, these labile exudates are metabolized by the active rhizosphere microbial communities (Waldrop & Firestone, 2006).

We found that total ecosystem ^13^C assimilated (above + belowground ^13^C content) at the 4-week sampling time was statistically indistinguishable from the belowground assimilated ^13^C at the 6-month sampling time. Root growth of annual species characteristic of these systems has been shown to decline by March (Jackson et al., 1989), suggesting that the ^13^C recovered in the belowground pool at the 4-week sampling time may have been primarily composed of structural plant root compounds. Additionally, under typical rainfall conditions, soil respiration in California annual grasslands has been shown to greatly decline by early April (Eviner, 2001), as sources of labile C substrates dwindle, and soil moisture begins to decline. This could explain why we saw little evidence of late growing-season decomposition or loss of ^13^C. ^13^C recovery in the belowground pool at our 6-month sampling time was 64% of the total ^13^C present immediately following the labeling period. Other studies in California annual grassland ecosystems have described similarly high accumulations of C (Schaeffer et al., 2017), and added ^13^C (Castanha et al., 2018) over the summer dry period. This appears to occur because of the large C input as dead root litter (following annual plant senescence and death in June-July each year), and the reduced ability of microbes to access and decompose this C due to the very low soil moisture characteristic of the Mediterranean-type summer; desiccation results in very low activity of decomposers as well as physical isolation from C substrates (Blankinship & Schimel, 2018).

### 4.2 Soil density fractions

In our study, the vast majority of soil organic carbon was recovered in the HF pool. We assume the heavy fraction is primarily composed of mineral-associated organic matter (MAOM), i.e. organic matter stabilized via mineral sorption and co-precipitation mechanisms (Kogel-Knabner et al., 2008). Mineral-associated SOM generally has a C:N ratio that aligns with the C:N ratio of microbes (∼10), as opposed to living plant material (>20), (Clemente et al., 2011). The C:N ratio of the HF in our study was consistently near 9:1, suggesting this HF carbon is largely of microbial origin. It has been shown that microbial products are more efficiently and effectively adsorbed to and thus stabilized by soil minerals than are plant-derived compounds (Lavallee et al., 2019). Furthermore, microbial transformation of detrital carbon inputs has been shown to be a critical precursor to long-term carbon stabilization (Cotrufo et al., 2013).

The C:N ratios of the FLF and OLF were approximately 20:1 and 17.5:1 respectively, reflecting a more plant-like signature than the HF material, and the C:N ratio of the OLF was consistently slightly lower than that of the FLF. As well, we found slightly higher total carbon in the OLF than in the FLF. California annual grasslands are typically dominated by annual herbaceous plant communities whose litter decomposes completely within three years (Eviner & Firestone, 2007), resulting in minimal FLF accumulation compared to other ecosystems (Crow et al., 2007). C-rich fungal hyphae, bacterial EPS, and plant mucilage promote soil aggregation both as physical structuring agents as well as major contributors to aggregate-associated soil carbon (Six et al. 2004). OLF material has been found to be largely dominated by fine root fragments and fungal hyphae (Kakouridis et al., 2021). The intermediate OLF C:N ratio we measured (between that of FLF and HF) suggests that microbial constituents and aggregation mechanisms such as fungal hyphae and bacterial EPS may have contributed to OLF formation in our samples, although extensive microbial processing of OLF material is likely limited due to physical protection mechanisms. If we interpret C:N ratios as an indicator of decomposition, then our data suggests increasing decomposition progressing from FLF to OLF to HF (Hyvonen et al., 1996).

The source compounds of ^13^C recovered in the density fractions likely differs by season. ^13^C recovered from Spring18-harvested soil likely represent mostly rhizodeposits and labile carbon substrates exuded by roots and possibly consumed by root-associated bacteria and fungi as well as AMF-mediated carbon flow from ^13^CO_2_-labeled plants (Kakouridis et al., 2021). By our Fall18 sampling, which occurred at the end of the summer dry period, much of the ^13^C in FLF would have been composed of senesced/dead ^13^CO_2_-labeled root detritus; we assume that most of the ^13^C in the HF and OLF was root-derived (Jackson et al., 2017). The onset of the 2018-2019 growing season (due to fall and winter rainfall), likely triggered decomposition of this senesced ^13^C-labeled shoot material derived from the previous growing season. Some of this decomposing ^13^C-labeled material could have been incorporated into the soil, representing an additional potential ^13^C source by the Spring19 sampling time. By Spring20 we assume that ^13^C recovered in the soil density fractions largely represents soil organic carbon that persisted from Spring19.

^13^C-SOC was recovered in all three density fractions at 4 weeks, 6 months, 1 year, and 2 years after ^13^C labeling (Figure 3). We observed a conspicuous increase in FLF ^13^C content between 4-weeks and 6-months sampling times due to rapid incorporation of ^13^C-labeled root detritus into this fraction following plant senescence and death, as well as likely incorporation of some aboveground litter; however, an equally conspicuous decrease in ^13^C within this fraction was observed between 6 months and 1 year after ^13^C labeling. Between the Fall of 2018 and Spring of 2019, about 38% of FLF material was either converted into more protected forms or lost as CO_2_. Such seasonal transience was not as apparent in the OLF or HF.

Additionally, of the three fractions, ^13^C enrichment was generally lowest in the OLF, indicating that this fraction is not as dynamic as the FLF, or perhaps even as the HF. This could suggest that the incorporation of “new” carbon into soil aggregates occurs less rapidly than does association of this carbon with minerals, or that stable aggregate formation requires repeated iterations of certain environmental conditions, such as annual plant growth periods or seasonal wet-dry cycles, the extent of which were not captured in the timespan of this study (Totsche et al., 2018). However, over this 2-year study, we see the distribution of ^13^C among the density fractions approaching that of total C; it is likely that with additional time, the quantity of ^13^C remaining in the OLF will exceed that remaining in the FLF (Figure 3).

The rapid association of rhizosphere-derived carbon with mineral surfaces was also observed by *Neurath et al. 2021,* in similar soils, who characterized the short-term dynamics (3 months) of root-input carbon. In that study, over the course of a 2-month incubation, the flux of new carbon onto and off of the mineral surfaces was substantial (accounting for over 6% of total C) while total mineral-associated C remained constant. For our study, this finding implies that some of the HF association with ^13^C OM could in fact be more dynamic than what we observed. By our two-year sampling, 77% of initial ^13^C stock had been lost from the system. However, of the carbon remaining, over 50% was in a protected form (occluded within soil microaggregates or associated with soil minerals) (Table 2). Furthermore, by four weeks after ^13^C labeling, we recovered 61% of soil ^13^C in the HF (Table 2), and quantity of ^13^C recovered in HF material did not detectably decline over the course of the study. This observation supports the importance of root and rhizosphere-derived carbon inputs in SOM formation (Sokol & Bradford, 2019; Pett-Ridge & Firestone, 2017), and is consistent with rapid microbially-mediated stabilization of carbon onto mineral surfaces (Kallenbach et al., 2016).

Our work indicates that initial associations between minerals and root-derived organic matter are significant and form rapidly. While the majority of annual C inputs to soil cycle rapidly (<2-year timescale), a sizeable proportion (11.5% original ^13^C inputs persist in HF by year 2) can potentially persist longer. In a study modeling soil carbon turnover in a California grassland site with similar physical characteristics to ours, Torn et al. (2013) calculated that 7% of HF carbon sampled from the 1-15cm depth was “fast” cycling (<2 year turnover) and 93% cycled on a centennial timescale. Our observations provide support for the existence of this small yet rapidly cycling HF carbon pool, but also suggest that the HF does not represent a single C pool operating on a single timescale, but rather multiple pools operating on multiple timescales (Lehmann & Kleber, 2015).

### 4.3 ^13^C NMR on heavy fraction

The combination of techniques used in this study provides a window into the dynamics and chemical characteristics of MAOM formation. The labeling of field plots with ^13^CO_2_ with subsequent sampling and soil density fractionation allowed us to trace the flow and fate of plant-derived inputs into organo-mineral associations. By applying ^13^C CPMAS NMR spectroscopy to our ^13^C labeled heavy fraction samples, we sought to further resolve the chemistry of “new” plant-derived carbon in the heavy fraction.

It is generally accepted that microbial products are a dominant source of mineral-stabilized organic matter that builds up in heavy fraction material over time (Preston et al., 2009; Creamer et al., 2019). The alkyl C functional group may represent a variety of microbial and plant-derived aliphatic compounds such as lipids, proteins, and waxes (Kögel-Knabner, 1997). In fact, in HF material isolated from a similar site at HREC, Neurath et al. found mineral-associated lipids were largely microbially derived (Neurath et al., 2021). Peaks in the O-Alkyl C region may represent various carbohydrates, proteins, and amino acids (Mathers et al., 2007), including plant-derived carbohydrates such as cellulose and hemicellulose (Kögel-Knabner, 1997), as well as N-rich proteins and root-exudate derived sugars (Angst et al., 2018). However, O-Alkyl C proportions in HF samples have also been attributed to carbohydrates of microbial origin (Schöning et al., 2006). The aromatic C functional group can contain plant-derived phenolic compounds such as lignin and tannins, aromatic portions of proteins and amino acids, as well as condensed, chemically resistant “black carbon” derived from historically frequent wildfires that occurred in California annual grasslands (Sanderman et al., 2008; Czimczik & Masiello, 2007). The carboxyl C functional group has been described to encompass highly oxidized C forms such as organic acids, ketones, and aldehydes (Mathers et al., 2007), and can be used as an indicator of microbial processing (Ng et al., 2014).

^13^C NMR spectra are only sensitive to molecules containing ^13^C. In Spring, 2018, we introduced ^13^C into the soil. The NMR spectra of the mineral-associated samples collected over the following two years show a decline in the proportion of ^13^C in the alkyl C functional group, and an increase in the proportion of ^13^C in the carboxyl C, aromatic C, and carboxyl C functional groups. The relatively constant quantity of ^13^C present in the HF over the course of the study suggests that alkyl ^13^C is being converted to carboxyl ^13^C, or that perhaps, that a portion of HF ^13^C is transient as suggested by Neurath et al., (2021), and that over the course of our study, some alkyl ^13^C is lost and some carboxyl ^13^C is accumulated. Carboxyl C content can be indicative of highly oxidized organic matter (Kogel-Knabner et al., 2008). As such, the increase in the relative proportion of carboxyl C we see over the 2-year study period could have resulted from the oxidation of organic matter during the process of decomposition (Baldock et al., 1992). The fact that we see a higher proportion of alkyl C shortly after the ^13^CO_2_ labeling period than we do two years later suggests that rhizodeposit C is a substantial source of rapidly forming mineral-alkyl C associations, whether as plant waxes such as cutin, or microbial lipids as suggested by Neurath et al., (2021).

Within a two-year period, we see an evolution of the effect of newly introduced ^13^C on the ^13^C NMR spectra of the heavy fraction (Figure 4). We expect that some of the added ^13^C in this fraction underwent chemical transformations during this period and that over time, only the most persistent ^13^C-HF associations will be retained as less persistent associations disappear. This could leave a legacy of the added ^13^C label that persists for decades or even centuries (Baisden et al., 2002), while the overall effect of the added ^13^C label should decline over time.

## 5. Conclusion and future directions

Understanding the patterns and control of soil organic carbon cycling in California annual grasslands is a critical precursor to any soil carbon management efforts within these ecosystems. Our study traced soil organic carbon formation from plant photosynthesis through its movement into and between soil fractions—and chemical forms in the heavy fraction—for two years. By applying a combination of analytical and spectroscopic techniques, we followed plant carbon from living roots to root detritus to occluded carbon and mineral-associated pools. About a quarter of the C fixed during the 5-day labeling was still present two years after the labeling; most of that “2-year old” carbon was found in the mineral associated heavy fraction. Solid-state ^13^C NMR spectroscopy was sufficiently sensitive to the ^13^C introduced that we were able to detect the photosynthetically derived carbon movement into and through the components of the mineral-associated carbon pool. This “new” carbon appears to have a distinct chemical fingerprint from the total background C; that spectroscopic profile declines over the 2 years in the field.

^13^C analysis and ^13^C NMR both revealed that the movement of carbon into the heavy fraction occurred rapidly in our system, within four weeks of plant photosynthesis, and that the influence of this “new” carbon on the total background C persisted over the course of our study. Future research that distinguishes between the labile vs. litter inputs on stabilized soil organic carbon formation, and that tracks those dynamics over decadal timescales, would further resolve how plant community characteristics (plant growth stage, lifecycle, seasonal climate parameters) influence the accrual and persistence of soil organic carbon and help us to better predict the responsiveness of annual grassland ecosystems to soil carbon management.

## Acknowledgements

This research was supported by the US Department of Energy (DOE) Office of Science, Office of Biological and Environmental Research Genomic Science program under award DE-SC0016247 (to MKF) and awards SCW1589, SCW1421 and the LLNL Soil Microbiome SFA, SCW1632 (to JPR). Work conducted at Lawrence Livermore National Laboratory was supported under the auspices of the U.S. DOE under Contract DE-AC52-07NA27344. Work conducted at Lawrence Berkeley National Laboratory was supported under Contract DE-AC02-05CH11231. Soil and plant collection and field plot management was supported by the Hopland Research and Extension Center.

---

**Supplemental Figure 1.**
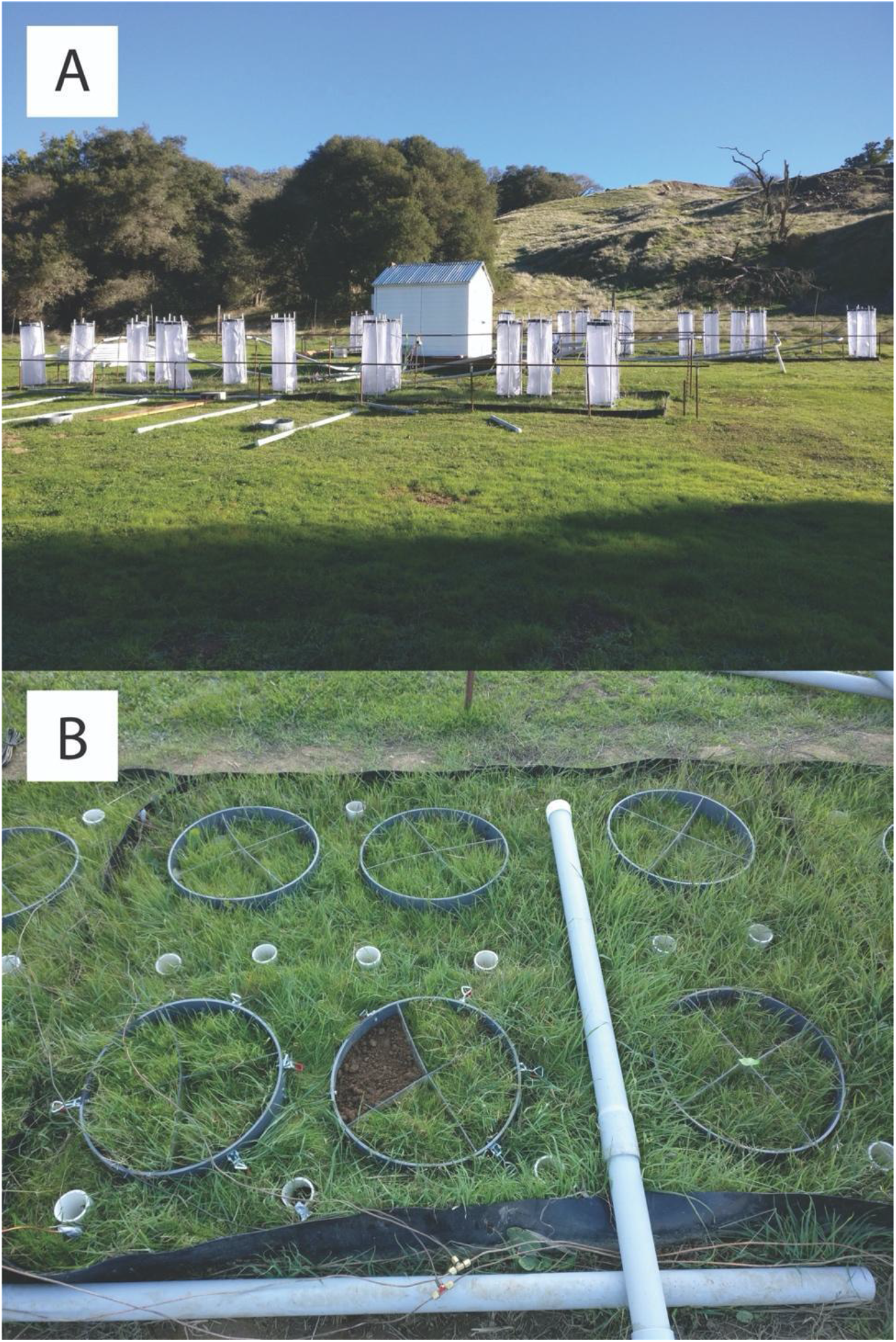
Field site ^13^CO_2_ labeling and sampling design. (A) ^13^CO_2_ labeling chambers, (B) Field plot with 6 circular subplots; subplots divided into 4 “wedges” for destructive sampling—ex. Bottom row center.

**Supplemental Figure 2.**
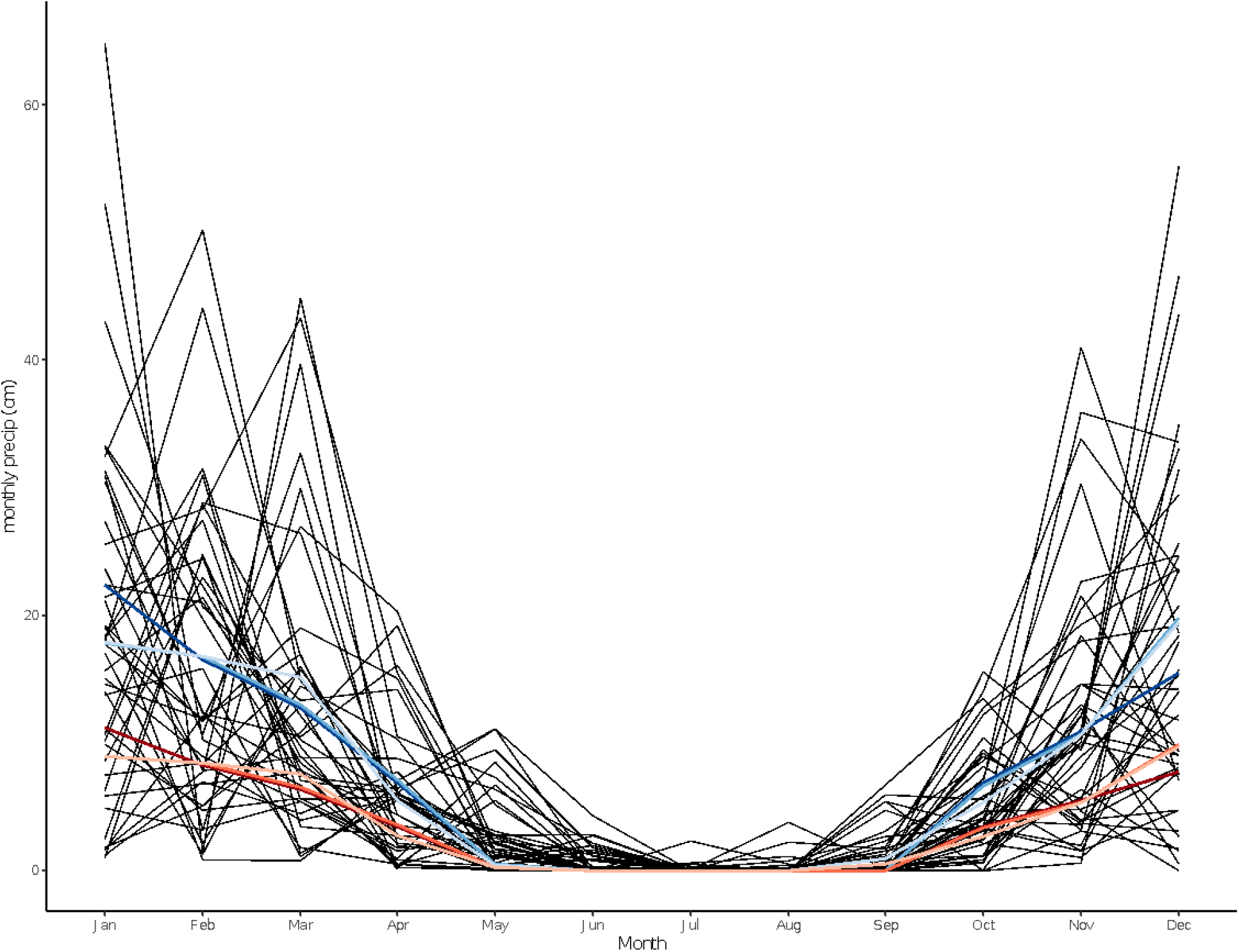
50-year annual precipitation + Annual rainfall for 50% and 100% precipitation treatment plots for 2017-2018, 2018-2019, and 2019-2020 growing seasons. Black lines indicate monthly precipitation at Hopland Research and Extension Center (HREC), our study site, dating back to 1970. Blue lines indicate manipulated rainfall received by our 100% of average annual precipitation treatment plots and red lines indicate manipulated rainfall received by our 50% of average annual precipitation treatment plots. Color shade scales from dark light with darkest blues and reds corresponding to the 2017-2018 growing season, and lightest shades corresponding to the 2019-2020 growing season.

**Supplement Figure 3.**
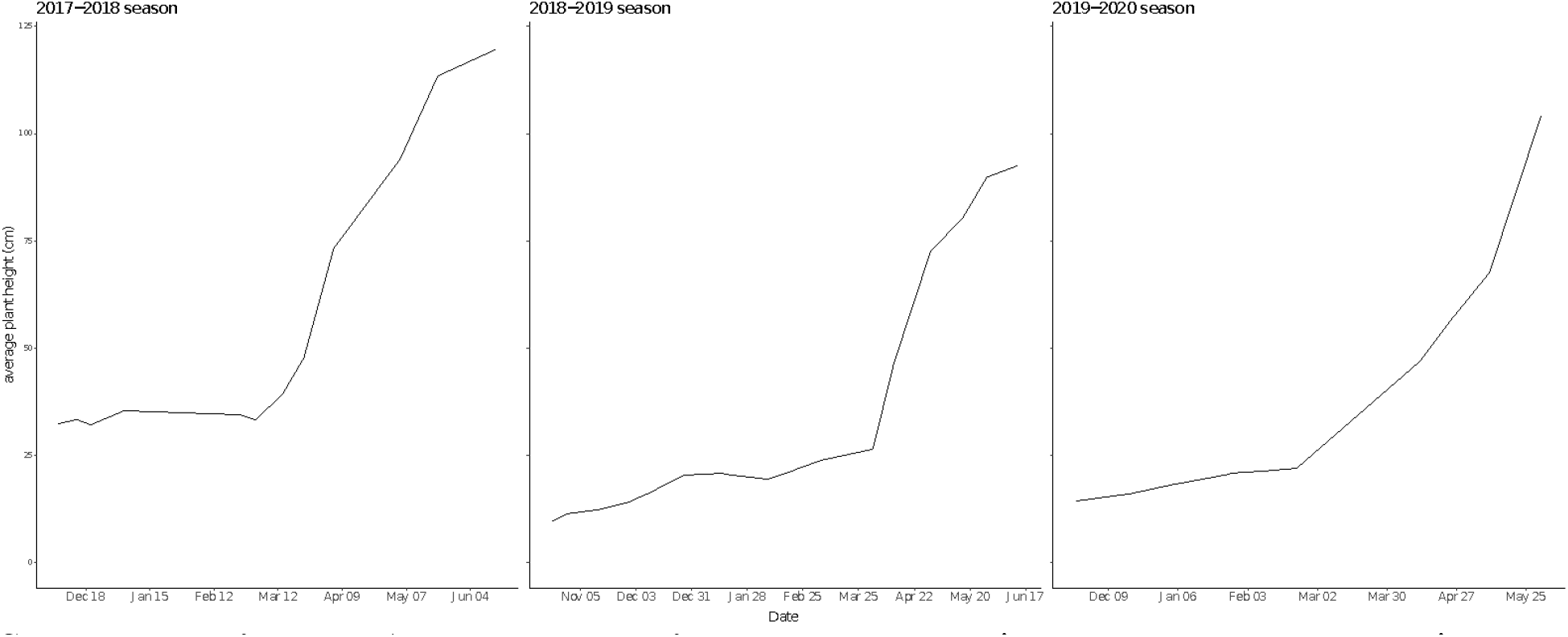
Average plant heights per season. Lines represent average maximum plant height throughout the three growing seasons. Maximum plant height was measured for each plot (n=16) and was measured in cm.

**Supplemental Figure 4.**
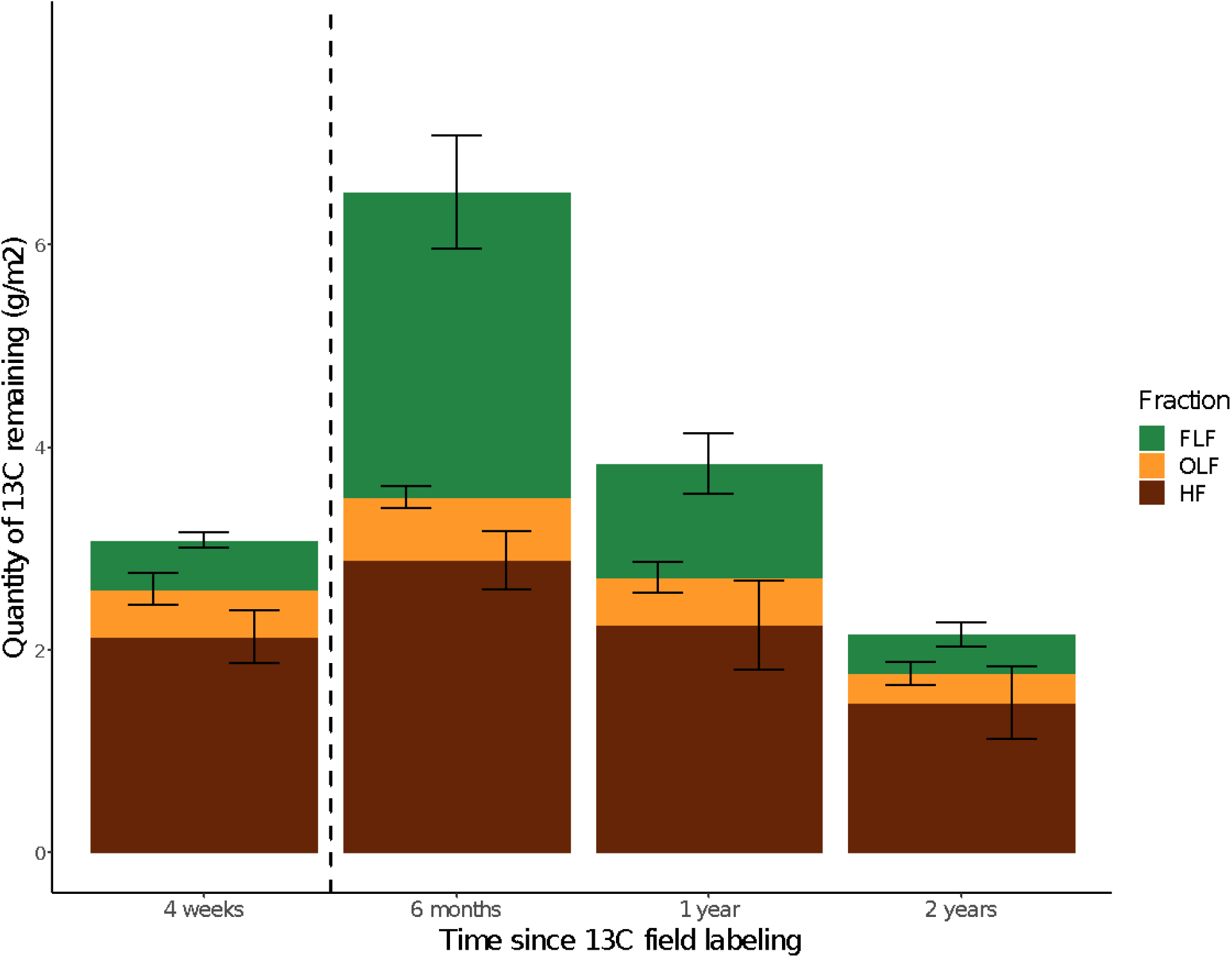
^13^C assimilation among soil density fractions. Total excess ^13^C content of soil density fractions: free-light fraction (FLF); occluded-light fraction (OLF) and heavy fraction (HF), scaled to soil volume (15cm sampling depth). Error bars represent 1 SE.

**Supplemental Figure 5.**
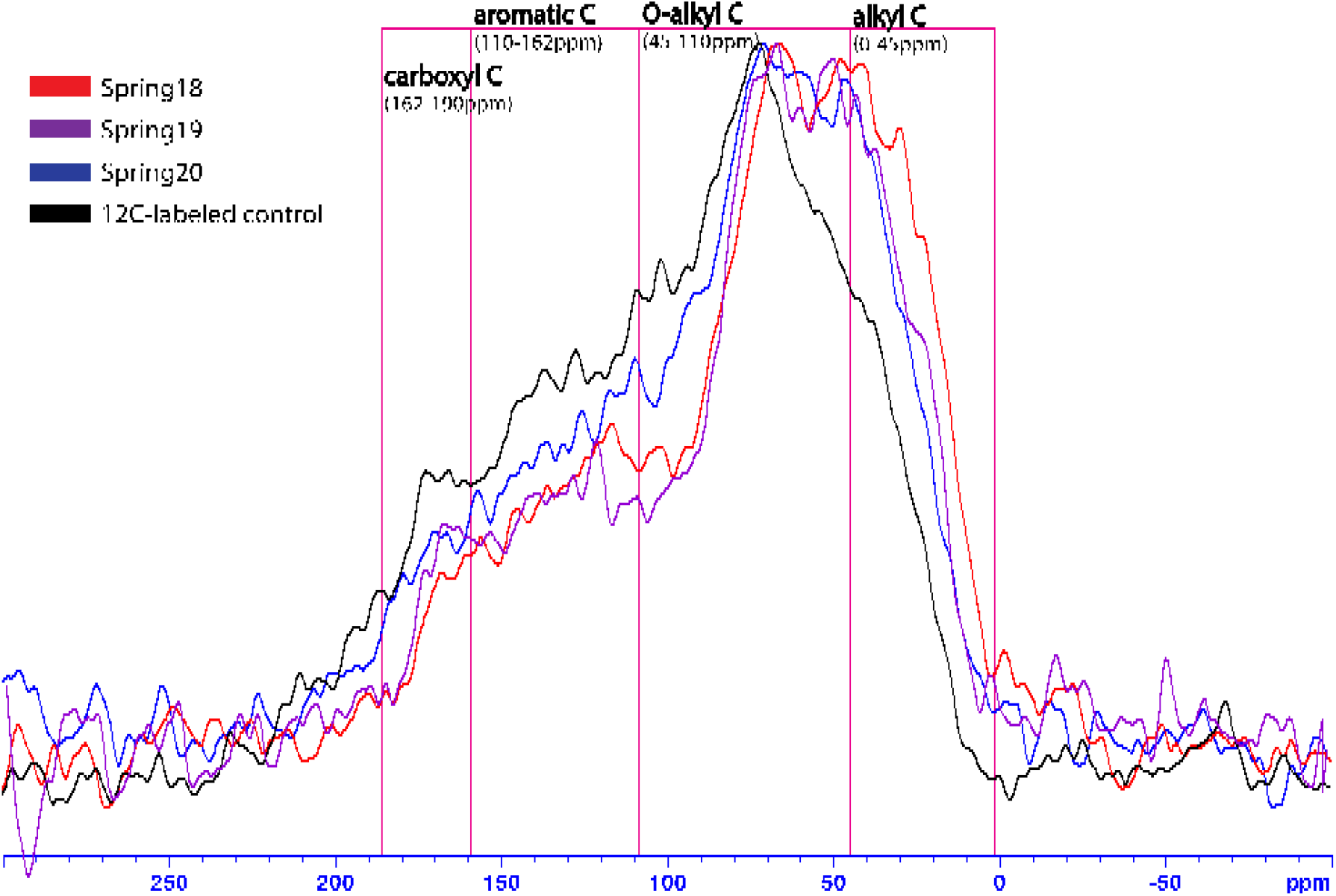
^13^C CPMAS NMR Spectra on Heavy Fraction 4 weeks, 1 year, and 2 years after ^13^C labeling relative to control sample. ^13^C NMR spectra of heavy fraction soil collected in Spring18 (4 weeks after ^13^C field labeling), Spring19 (1 year after), Spring20 (2 years after), and a ^12^C-labeleled control (collected in Spring18). Carbon functional groups outlined in pink.

**Supplemental Figure 6.**
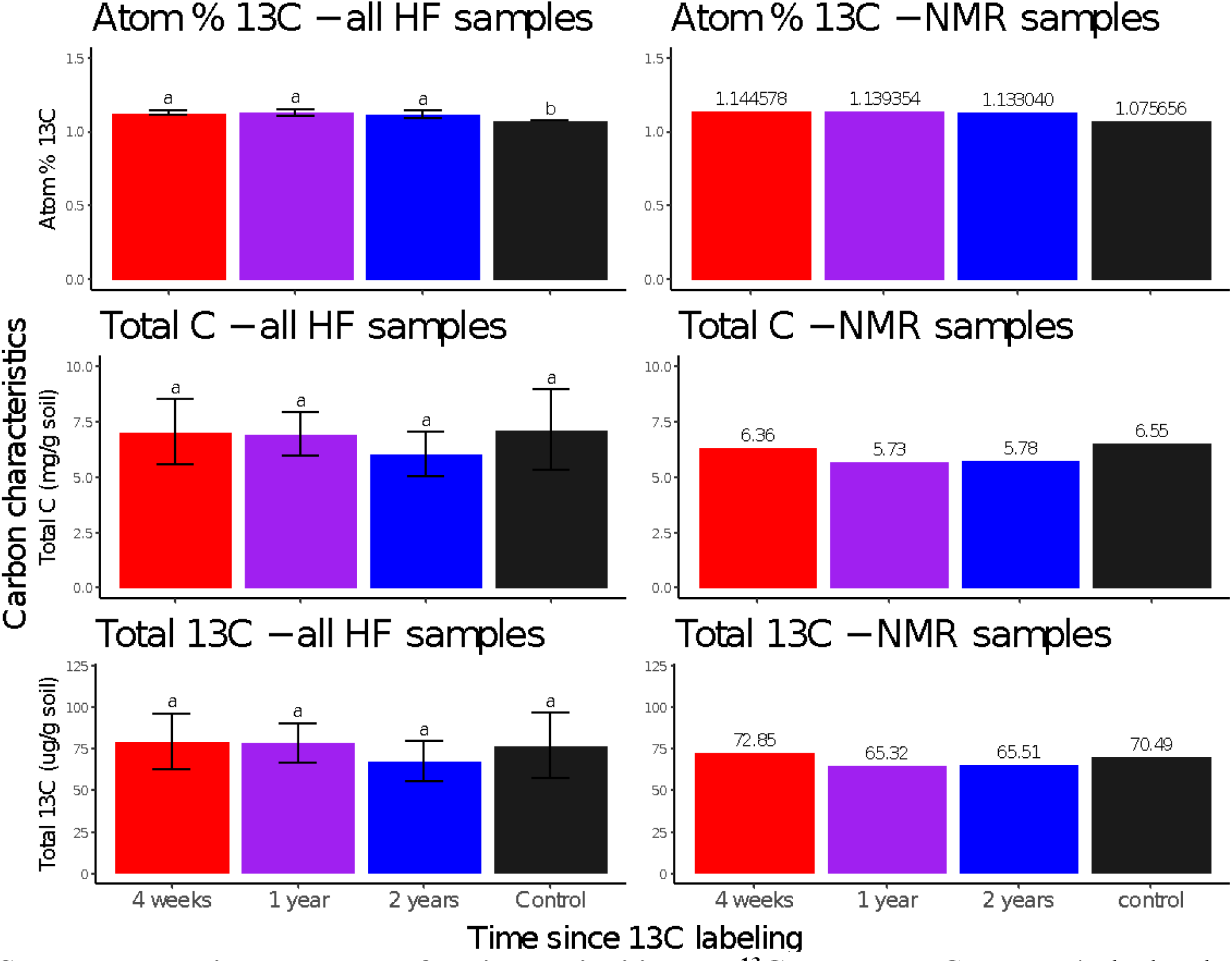
Heavy fraction variability and ^13^C content. 13C content (calculated from total C content X atom-% ^13^C) measured for all 8 HF replicates of individual samples for which ^13^C NMR spectra were acquired (left column), and for individual samples for which ^13^C NMR spectra were acquired (right column). Error bars represent 1 SD. Significant differences by Tukey’s HSD (p-value = 0.05) between timepoints are indicated by letters. Atom-% ^13^C, total C, and total ^13^C values for the HF samples for which ^13^C NMR spectra were acquired (right column) label each bar. Colors represent timepoints and correspond to spectra represented in Supplemental Figure 2.

